# Practical considerations in the use of a porcine model (*Sus scrofa domesticus*) to assess prevention of postoperative peritubal adhesions

**DOI:** 10.1101/675074

**Authors:** Claudio Peixoto Crispi, Claudio Peixoto Crispi, Fernando Luis Fernandes Mendes, Claudio Moura de Andrade, Leon Cardeman, Nilton de Nadai Filho, Elyzabeth Avvad Portari, Marlon de Freitas Fonseca

## Abstract

Infertility has been a common postoperative problem caused by peritoneal adhesions. Since several prophylactic agents have recently shown promising preliminary results, more complete studies comparing their real efficacy and safety are needed urgently. The aim of this study was to investigate and describe practical considerations of a porcine model that can be used to assess such prophylactic agents. First, 10 healthy 5½ months old female pigs (24.3 – 31.3 Kg) underwent a standardized laparoscopy to provoke peritubal adhesion formation without prophylactic agents. After 30 days, a second-look laparoscopy was performed to evaluate adhesions and perform adnexectomy for histopathological evaluation. Adhesions at different sites were classified by grade, for which the scores range from 0 (no adhesion) to 3 (very strong vascularized adhesions), and also by area, with scores ranging from 0 (no adhesion) to 4 (>75% of the injured area). The histopathological evaluation of the distal uterine horns, oviducts and ovaries were compared withthose from a control group of six healthy pigs with no previous surgery. Biological samples were collected to assess vitality, inflammation and renal, hepatic and hematopoietic systems. There were small (but significant) changes in serum albumin (P=0.07), globulin (P=0.07), C-reactive protein (P=0.011), fibrinogen (P=0.023) and bilirubin (P<0.01) after 30 days, but all values were within the normal range. No inflammation or abscess formation was observed, but different degrees of adhesion were identified. The estimated occurrence of adhesion (scores >0) and of strong / very strong adhesion (scores >1) was 75% (95% CI: 55 – 94.9) and 65% (95% CI: 45 – 85), respectively. The porcine model represents a useful animal platform that can be used to test the efficacy and safety of candidate prophylactic agents intended to prevent postoperative peritubal adhesions formation. We present several practical considerations and measures that can help to minimize animal suffering and avoid problems during such experiments.

## Introduction

Abdominal intraperitoneal postoperative adhesions are fibrous bands that span two or more organs or the inner abdominal wall. Such adhesions usually develop as a consequence of the healing process in peritoneum that was injured during surgery, regardless of the surgical approach [1].

Perioperative and postoperative complications of adhesions include accidental abdominal viscera injuries (when a new laparoscopic puncture is made), longer duration of subsequent surgeries, chronic abdominopelvic pain, and intestinal obstruction [2]. In women, adhesions also can impair fertility by distorting adnexal anatomy and interfering with gamete and embryo transport [3]. As a result of concern about such complications, the number of publications about surgical adhesions has grown year after year, with considerable interest in recent years in novel prophylactic agents and methods that have shown intriguing promise [4,5,6,7,8,9].

Although body mass index (BMI) and several preoperative inflammatory blood biomarkers have emerged as potential predictors of post-operative abdominal adhesion formation [2], recent reviews emphasize their shortcomings and affirm that optimal approaches to adhesion formation prevention still elude us. Several articles have suggested priorities for future research. Future studies, these authors say, should consider how adhesion prophylaxis can preserve fertility, include assessments of the safety of the prophylactic agents, assess adhesions in a uniform or standardized way, and present complete statistical analyses [10,11]. The authors also call for non-industry funding so that the research is untainted by pharmaceutical manufacturers’ financial support [10,12].

Since more complete studies assessing the effectiveness and safety of adhesion prophylactic agents are necessary [9,13], the search for a safe, efficient, and easy-to-use method of adhesion prophylaxis starts by determining a qualified animal model [7]. Before assessing prophylactic agents and methods in humans, controlled experiments to evaluate the efficacy of prophylactic products in animal models have usually followed a simple methodology. First, a standardized surgical injury (able to provoke adhesion formation) is performed and, after randomization, the tested agent or a control substance is applied. Then, after a period of time long enough to form adhesions, the two groups are compared in relation to adhesion formation. Although most animal studies of postsurgical adhesions have used small animals (i.e. rats and rabbits) due to practical considerations [13], some research groups have elected to use porcine models because of their known efficiency – especially under laparoscopic conditions – and the potential to test prophylactic agents in more realistic conditions [14,15,16].

In this study, we sought to describe some major aspects of a porcine model used to assess postoperative peritubal adhesion formation, including the expected incidence of adhesions, the histopathological characteristics of the uterine horn, and the baseline values and natural changes in several biomarkers that are observed 30 days after a standardized peritubal tissue injury triggered by laparoscopy when no prophylactic agent was used.

## Materials and Methods

### Design and team

This experimental study was carried out through a partnership of three institutions: the Crispi Institute for Minimally Invasive Surgery (www.institutocrispi.com.br), the Suprema Faculty of Medical Sciences and Health of Juiz de Fora (www.suprema.edu.br), and the Research and Education Center for Phototherapy in Health Sciences (www.nupen.com.br). In order to establish a detailed protocol that also highlights various types of possible pitfalls or operational difficulties in future experiments, an interdisciplinary approach was used throughout the elaboration of this work. Thus, the planning and execution of this study included the participation of specialists from different areas: anesthesiology, gynecology, veterinary medicine, proctology, urology, clinical pathology, surgical nursing, quantitative methods and clinical laboratory analysis.

### Ethical statement

This experimental study was carried out at the Suprema Surgical Training Center (Juiz de Fora, Minas Gerais, Brazil) in strict accordance with the Guide for the Care and Use of Laboratory Animals of the National Research Ethics Commission of the Brazilian Ministry of Health and the recommendations of the National Centre for the Replacement, Refinement and Reduction of Animals in Research (London, United Kingdom). In order to maximize reproducibility and the potential for the re-use of data, we also followed the Animal Research: Reporting of *In Vivo* Experiments (ARRIVE) Guidelines [17].

The protocol was approved by the Institutional Animal Care and Use Committee (CEUA – Suprema; Protocol Number 004/2017). Besides the health certificate issued by a veterinarian provided by the supplier (Fazenda Penalva, Juiz de Fora, MG), the veterinarian responsible for the study (F.L.F.M.) clinically evaluated all the animals before and during this study.

In order to optimize the sample size, we considered as realistic an experimental model in which adhesion formation would occur in 90% rather than all of the animals [10,18,19,20]. Assuming a 20% error, and using the formula N = 1,96^2^ × P(1 – P) / D^2^, where N is the minimum sample size, P is the expected prevalence and D is the maximum accepted error, we calculated 8.6 as the minimum number of animals necessary to estimate the incidence of adhesions. Thus, this study included 10 animals.

### Animals and procedures

The study was carried out in two phases; the same two surgeons performed all surgeries. In the first phase, 10 healthy 5½ month-old female pigs (*Sus scrofa domesticus*; crossbreed Large White) underwent laparoscopy to execute standardized bilateral pelvic injuries in order to provoke the formation of peritubal adhesions. In the second phase, 30 days later, a “second-look” laparoscopy was performed to classify and quantify the peritoneal adhesions, and to perform adnexectomy for histopathological examination of the distal uterine horn, including ovaries and oviducts. The animals were then euthanized.

The animals, which had fasted for 12 hours, were premedicated with an admixture of midazolam (0.5mg/kg) + atropine (0.04 mg/kg) + ketamine (2 mg/Kg) + acepromazine (0.1 mg/Kg) – administered as a single intramuscular injection. General inhalation anesthesia was then induced with a swine-specific mask and maintained (after oral intubation) with isoflurane (1.5 – 2.5 vol.%) in oxygen (flow rate: 2 L/min). Monitoring instruments during anesthesia included pulse oximeter with plethysmograph, rectal thermometer and sphygmomanometer. The total perioperative hydration was standardized as intravenous infusion of 500 mL of sodium chloride 0.9%.

In the first phase (peritubal injury), the laparoscopic surgeries were performed in the evening between 5 pm and 11 pm on consecutive days (two campaigns). When the animal was adequately anesthetized, the abdominal region was scrubbed with warm water, shaved, and disinfected; thus the laparoscopic injury was performed in aseptic conditions. The ambient temperature of the operating suite was maintained between 21°C and 23°C. Immediately after orotracheal intubation, the veterinary anesthesiologist administered intramuscular Enrofloxacin 10% (2.5 mg/Kg) for antibiotic prophylaxis and Meloxicam 2% (0.4 mg/Kg) for preemptive analgesia. The pharmacological strategy for postoperative analgesia also included infiltration of a long-lasting local anesthetic into laparoscopic punctures at the end of surgery (detailed below).

The second-look surgery followed the same protocol, except for the postoperative analgesia, antibiotic prophylaxis and aseptic conditions.

### Laparoscopic protocol to form peritubal adhesions

Given the frequent difficulties and complications experienced using other techniques to perform the first puncture in pigs, our group considers and recommends the Veress needle technique as the best method to establish the pneumoperitoneum in these animals. In this study, after the first 11 mm trocar was inserted through a small incision in the umbilical scar, three accessory 5 mm trocars were inserted as illustrated in **Figure 1A**.

During laparoscopy, the animals were placed in Trendelenburg position, the carbon dioxide pressure was set in 10 mmHg with high flow insufflation for the maintenance of the pneumoperitoneum, and the surgery was performed as usually done in humans [22]. After a careful inventory of the entire abdominal and pelvic cavity in order to exclude naturally formed adhesions (**Figure 2A-B**), the first step of the injury was a laparoscopic suture in distal segment of the uterine horn with Polyglactin 910 (Coated Vicryl^®^ 2.0 ½ circle 31 mm, Ethicon) by introducing the needle into the broad ligament (**Figure 2C**) and performing a knot wrapping the entire circumference of uterine horn, similar to the Pomeroy technique (**Figure 2D**). Then, a small portion of the uterine horn (about 1 cm) was excised at a distal site close to the utero-tubal junction with a laparoscopic scissors (**Figure 2E-F**). Subsequently, in order to favor the peritubal adhesions formation, we performed bilateral excisions of an 8 cm × 10 cm area of the peritoneum of the pelvic sidewall located opposite the left and right uterine horns using both laparoscopic scissors and blunt dissection until the musculature was totally exposed (**Figure 2G-H**). Finally, a similar area of peritoneum was excised at the anterior wall below the umbilical scar, up to (but not reaching) the bladder. Rather than energy, only surgical gauze was used for hemostasis in all sites.

**Figure 1.**
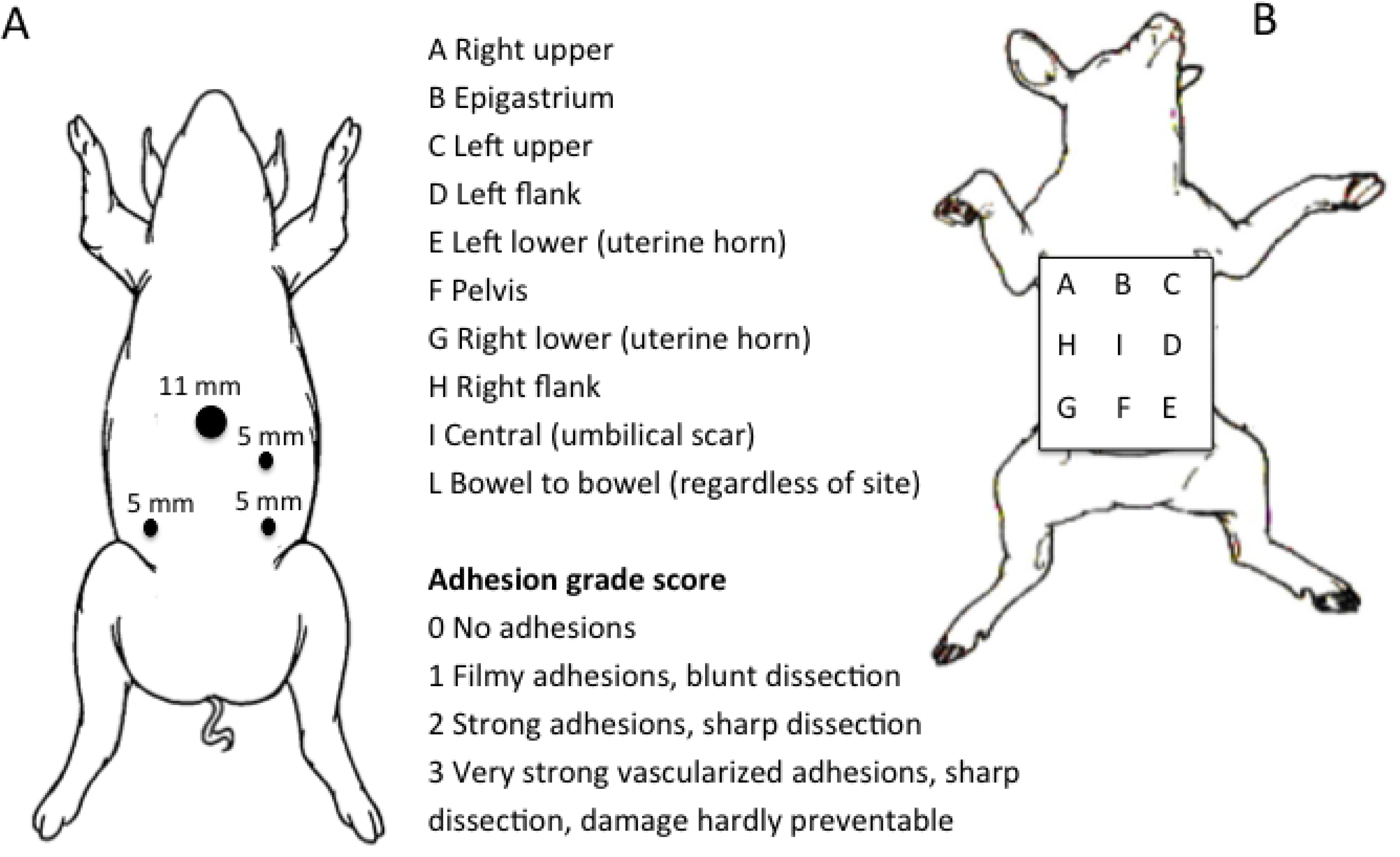
Schematic illustration of trocar placement in the porcine model (**A**) and the regions of interest to calculate the Peritoneal Adhesion Index (**B**), which is the sum of the grade scores in all regions (Adapted from Coccolini et al., 2013) [21].

**Figure 2.**
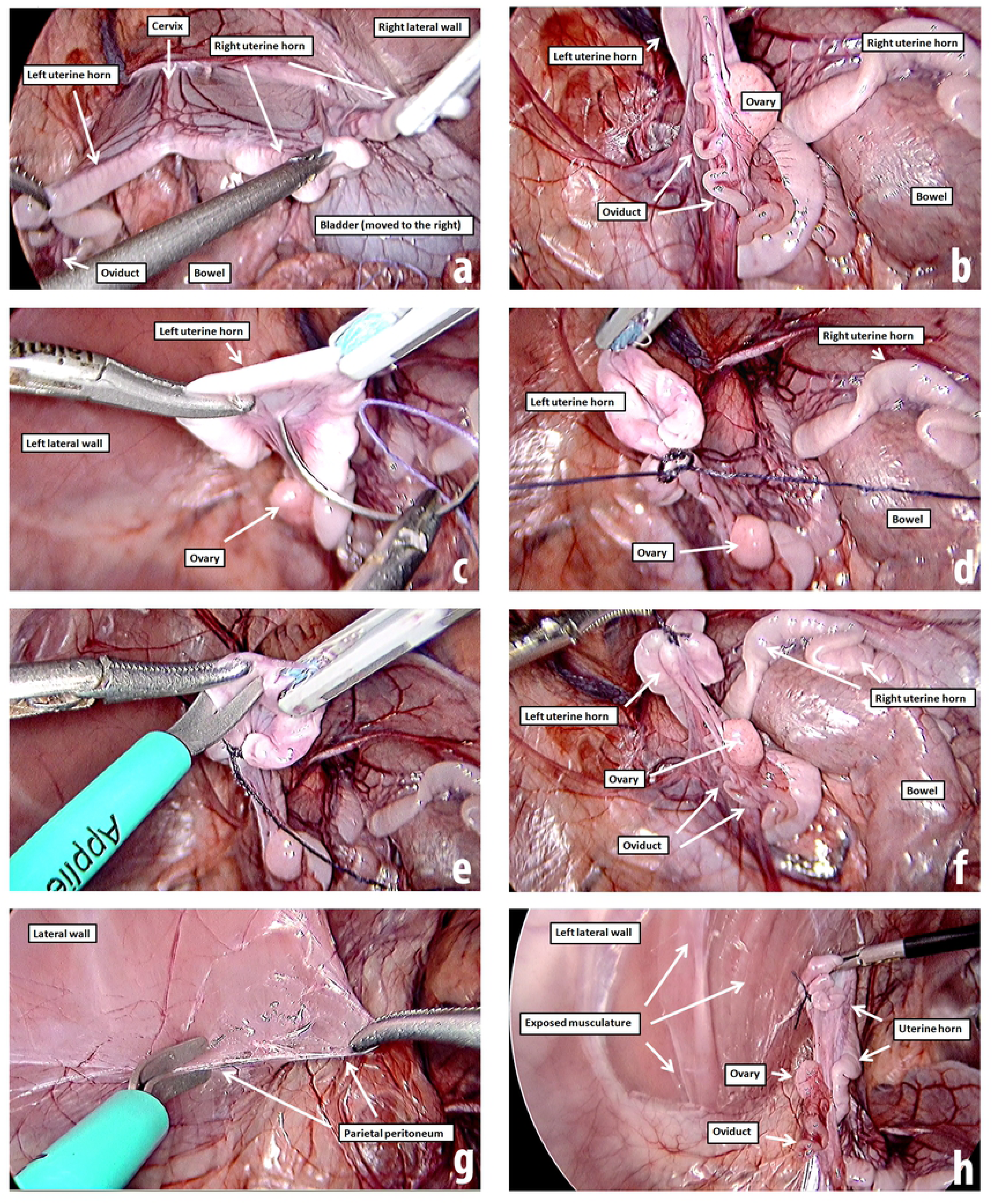
Laparoscopic protocol to form peritubal adhesions: panoramic view during inventory of the cavity (**A**); identification of the ovary and oviduct (**B**); laparoscopic suture in the uterine horn (**C**); knot wrapping the entire circumference of uterine horn (**D**); excision of a small portion of uterine horn (**E**); panoramic view of injured uterine horn (**F**); excision of about 80 cm^2^ of peritoneum on the pelvic sidewall (**G**); panoramic view at the end of peritubal injury (**H**). The protocol was performed bilaterally.

After the surgery was completed, in order to improve the postoperative analgesia [23,24], the surgeon injected 0.5% bupivacaine around each laparoscopic puncture: 2 mL in the site where the 11 mm trocar had been inserted, and 1 mL in the 3 other sites where the 5 mm trocars were placed. As young pigs have a relatively thin abdominal wall, these injections were made with a delicate needle (16 × 4.5 mm) in order to minimize the risk of drilling intra-abdominal structures.

### Postoperative care

After the first surgery (injury), the animals were allocated in groups of 3 or 4 animals in an infirmary housing, where they received care for 30 days in boxes of approximately 8 m^2^ with a fenestrated bottom, built specifically for this purpose. The animals were fed a special ration for pigs (500 g / day / animal during the first 2 weeks; 600 g / day / animal afterwards) and oral hydration (*ad libitum*) by an automatic system with water from the public network. After surgery, a veterinarian examined the animals three times a day during the first week, and the basic care included not only a veterinary topical antiseptic, but also regular analgesia with intramuscular Meloxicam (4 days) and daily antibiotic prophylaxis with Enrofloxacin for one week. The animals were cared for by a caregiver under the supervision of the veterinarian until the “second-look” surgery. The main objective of this observance was to respond promptly to eventual clinical intercurrences and provide immediate diagnosis, specific treatment, and necropsy in case of death (none occurred). During the 30 days post-operative care, the temperature and humidity inside the housing was checked three times a day and ranged, respectively, between 16°C and 28°C (median 22°C) and between 60% and 99% (median 86%).

### Second look and assessment of peritubal adhesions

In this study, rather than necropsy, the presence of peritoneal adhesions in specific sites was assessed laparoscopically 30 days post-injury using a standardized classification and quantification methodology that is based on the macroscopic appearance of adhesions and their distribution in different regions of the abdomen (**Figure 1B**). The sites were classified using an ordinal variable (on a 0 to 3 scale) derived from the Peritoneal Adhesion Index [21], and also received a score (on a 0 to 4 scale) based on the ratio of the area of adhesion to the area of injury [13]. The excised area of the peritoneum (about 8 × 10 cm) was considered the reference area (**Figure 3**) to determine the adhesion area score (**Table 1**).

**Table 1.** Postoperative peritoneal adhesions (raw scores) assessed through a second-look laparoscopy performed 30 days after standardized bilateral tubal injury and excision of adjacent peritoneum of the pelvic sidewall toprovoke adhesion formation.

| Animal | 1 | 2 | 3 | 4 | 5 | 6 | 7 | 8 | 9 | 10 |
| --- | --- | --- | --- | --- | --- | --- | --- | --- | --- | --- |
| **Grade** |  |  |  |  |  |  |  |  |  |  |
| Left uterine horn site | 2 | 2 | 0 | 2 | 2 | 2 | 0 | 2 | 2 | 0 |
| Right uterine horn site | 2 | 2 | 2 | 2 | 0 | 2 | 0 | 1 | 1 | 2 |
| Bowel to bowel | 0 | 2 | 0 | 0 | 0 | 0 | 0 | 0 | 0 | 0 |
| PAI | 4 | 6 | 2 | 4 | 2 | 4 | 0 | 3 | 3 | 2 |
| **Area** |  |  |  |  |  |  |  |  |  |  |
| Left uterine horn site | 1 | 1 | 0 | 3 | 1 | 2 | 0 | 2 | 1 | 0 |
| Right uterine horn site | 1 | 1 | 1 | 1 | 0 | 1 | 0 | 1 | 1 | 1 |

**Figure 3.**
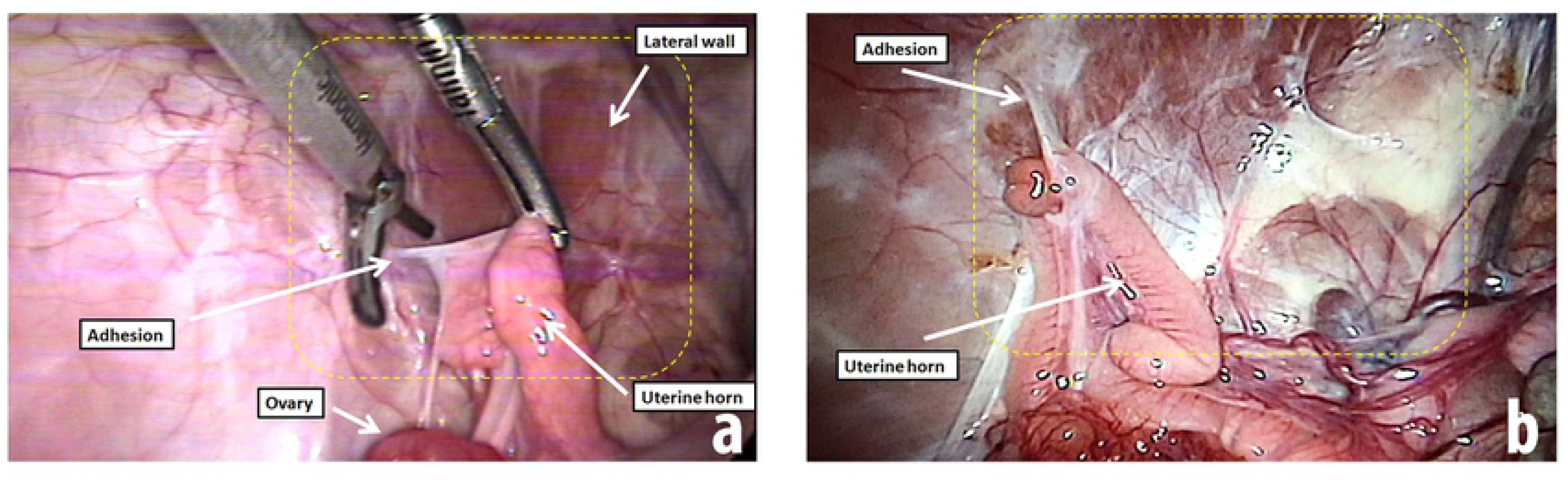
Laparoscopic view of peritubal adhesions in a young female pig during a second-look laparoscopy that was performed one month after a standardized laparoscopy to provoke adhesion formation. The yellow dotted line represents the peritoneum area that was removed from the lateral wall (approximately 80 cm^2^). Both images **a** and **b** exhibit adhesions that were classified as Peritoneal Adhesion Index raw score 2 (strong adhesions, sharp dissection) according to Coccolini et al., 2013 [21].

After evaluating adhesion scores under laparoscopic view, the distal uterine horn with oviduct (infundibulum, ampulla and isthmus) and ovary on each side were laparoscopically removed using an ultrasonic scalpel in order to minimize bleeding [25]. The harvested specimens were fixed in 10% neutral buffered formalin solution, and then embedded in paraffin, sectioned, and stained with hematoxylin-eosin for evaluation by a single experienced pathologist (L.C.). The histopathological assessment of the distal uterine horns, oviducts and ovaries from the ten 6½ month-old previously injured pigs were compared with those of a control group composed of six healthy 5½ months old pigs with no history of surgery, with a focus on the injury repair response and naturally occurring changes from 5½ to 6½ months.

Upon conclusion of the second look laparoscopy, all the animals were euthanized by deep anesthesia followed by intravenous administration of 10 mL of 19.1% potassium chloride.

### Sampling to assess toxicologic biomarkers

In order to explore the nutritional status, the immune response and the potential late consequences of surgery on the hematopoietic, renal and hepatic systems, blood samples were obtained at two moments: before the laparoscopic injury and before the second-look. The samples were collected systematically after orotracheal intubation by puncture of an animal’s ear vein with a 22G peripheral vein catheter while keeping the animal in dorsal recumbency. Because of difficulties encountered during preliminary blood specimen collections using Vacutainers, blood collection was performed by dripping blood from the open catheter directly into uncovered tubes. After filling each tube with the appropriate volume of blood, the tube was capped and gently shaken to provide contact with the anticoagulant avoiding coagulation. To ensure consistency, this maneuver was performed with the concurrent participation of two veterinarians.

Many disorders can be detected in their early stages by examination of the urine. Urinalysis was performed as a screening test to detect and/or measure by products of normal and abnormal metabolism (e.g. glucose, protein, bilirubin, red blood cells, white blood cells, crystals), and bacteria. Urine samples were systematically collected, but only during the second-look surgery due to concern about the risk of introducing infections while manipulating the urinary tract of young female pigs. In fact, catheterization of the bladder to collect the urine specimens proved so challenging, the specimens were obtained by transdermal suprapubic aspiration under laparoscopic vision using a 25 Gauge 3.5 inches Quincke spinal needle coupled to a 5 mL syringe. To favor diuresis and ensure 5 mL of urine could be collected at the end of the second-look surgery, bolus intravenous hydration with 500 mL of 0.9% NaCl solution was initiated following induction of anesthesia.

The clinical analysis laboratory responsible for the analysis of study biological specimens is accredited by the Brazilian Ministry of Agriculture, Livestock and Food Supply (Ministério da Agricultura, Pecuária e Abastecimento, MAPA) for the analysis of official samples. The laboratory has a quality control system that complies with NBR ISO 17025 standards; it assesses precision and accuracy daily, and undergoes external quality control.

## Statistics

The database was managed using Microsoft Office Excel^®^ version 2010 (Microsoft Corp., Redmond, WA, USA). Statistics and charts were generated using IBM^®^ SPSS^®^ Statistics Standard Grad Pack 20 (NY, USA). The statistical results were considered significant when P < 0.05 (2-sided).

## Results

### Time spent

The time spent with each animal during the first surgery ranged between limits that are considered acceptable. The median total time between the intramuscular administration of premedication and the induction of general anesthesia was 13 min (min 7, max 23 min); the median total time of general anesthesia (from intubation to extubation) was 79 min (min 68, max 104 min); the median total time of surgery (from the beginning of the first puncture to the last suture) was 50 min (min 32, max 71 min); and the median total time of pneumoperitoneum was 33 min (min 23, max 64 min).

### Peritubal adhesions after 30 days

No inflammation or abscess formation was observed in any of the operated animals whereas adhesions of varying degrees were identified (**Figure 3**). Due to its proximity to the pelvic sidewall, most of the adhesions that formed involved the uterine horn, as expected. Only one animal had an adhesion elsewhere: a bowel wall to bowel wall adhesion. When only the 20 uterine horn sites were considered, the estimated incidence of any adhesion with a score > 0 was 75% (95% confidence interval: 55 – 94.9) and the estimated incidence of a strong or very strong adhesion with a score > 1 was 65% (95% confidence interval: 45 – 85). The raw scores characterizing the postoperative peritoneal adhesions are presented in **Table 1**.

PAI (Peritoneal Adhesion Index) is the sum of the raw scores in all regions (Adapted from Coccolini et al., 2013) [21]: 0 No adhesions; 1 Filmy adhesions, blunt dissection; 2 Strong adhesions, sharp dissection; 3 Very strong vascularized adhesions, sharp dissection, damage hardly preventable. Area scores (adhesion area / injured area ratio): 0 (no adhesion); 1 (≤25% of initial injured area); 2 (>25% and <50% of initial injured area); 3 (50% – 75% of initial injured area) or 4 (>75% of initial injured area) [13]. The excised area of the sidewall peritoneum adjacent to each uterine horn was considered as a reference injured area (about 8 × 10 cm). Sites with no adhesions in all ten animals (all scores = 0) were not included in this table. The animals were numbered in order according the time of the first surgery (injury).

### Histopathological observations

Despite being somewhat heterogeneous, the histological findings in all assessed uterine horns (including the oviduct, ovary and uterus) may be considered consistent with an inflammatory response during the natural healing process evoked by the injuries performed 30 days earlier.

In swine, as in other mammalians, the histology of the ovary – both its outer cortex and inner medulla – varies with the age and phase of the sexual cycle. The surface of the pigs’ ovaries is covered by a low cuboidal epithelium and, immediately beneath this surface epithelium; there is a dense connective tissue sheath, a tunica albuginea. The cortex is composed of ovarian follicles, which usually occur in different stages of development (least mature to most mature). The inner medulla is composed by a loose connective tissue that contains nerves, blood vessels and lymph vessels which enter the ovary at the hilus from the mesovarium.

In the pilot study assessing 5½ month-old pigs with no previous surgery (n=6; control group), we observed in 5 of them that the cortex was mainly disrupted by numerous primordial follicles, primary and secondary follicles. In only one case, there was also tertiary or cystic follicles that are usually located at the periphery. In this series, nineteen ovaries from the ten 6½ month-old pigs that were injured 30 days earlier showed a lobulated surface – “blister like” structures – easily visible to the naked eye. When the ovaries were cut, the general macroscopic aspect was multicystic measuring from 1 to 5 mm in diameter. Histologically, the ovaries presented many tertiary or cystic follicles amid the least mature follicles. In three of these twenty ovaries, fibrosis was observed in connective tissue from the mesovarium (**Figure 4**).

**Figure 4.**
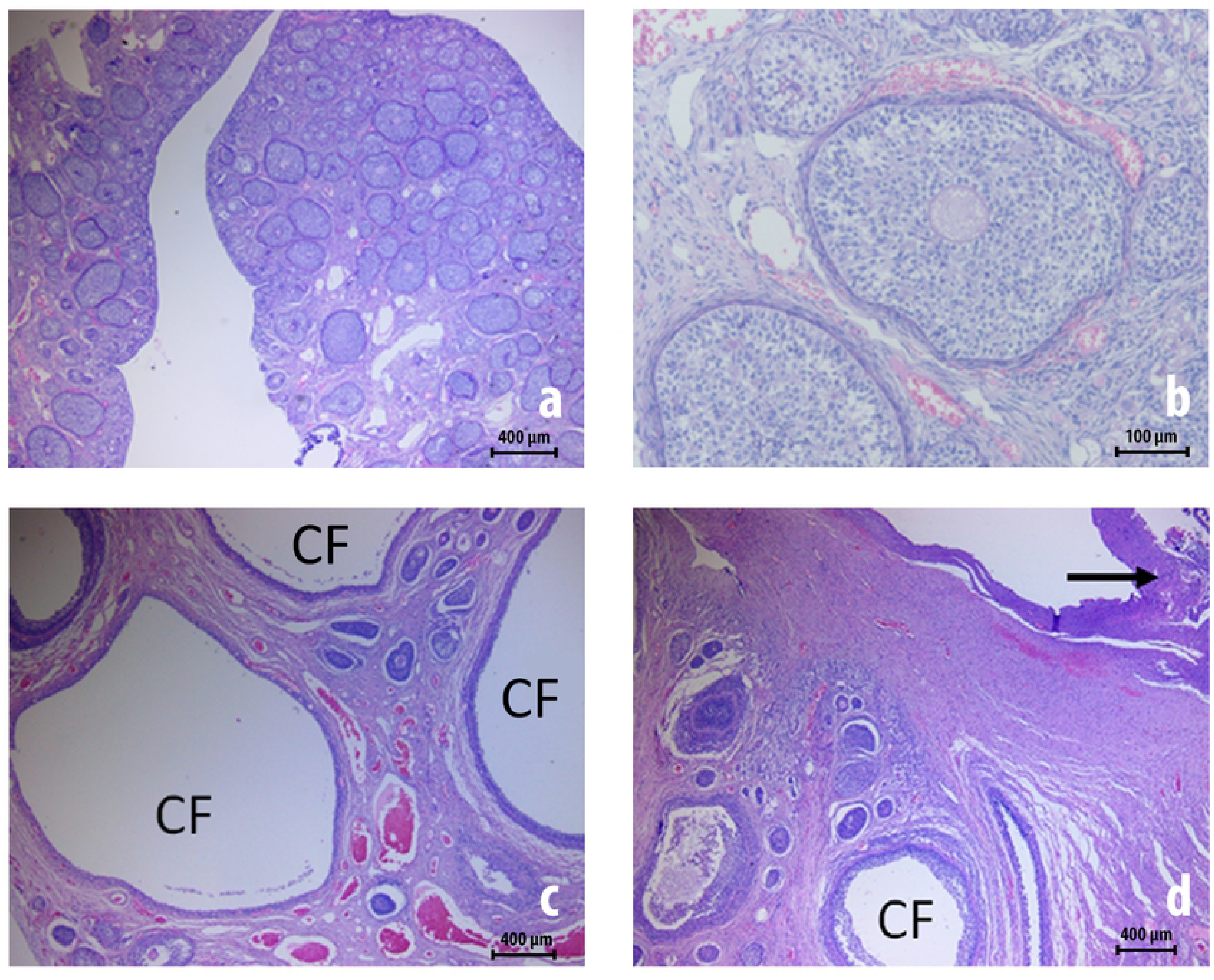
Ovarian histology prior to and 30 days after peritubal injury (hematoxylin-eosin staining). Images **a** and **b** exhibit micrographs of an ovary from a 5½ month-old pig with no previous surgery (control group) showing ovarian follicles in different stages (primordial, primary and secondary follicles). Images **c** and **d** exhibit an ovary from a 6½ months old pig assessed 30 days after a standardized peritubal laparoscopic injury, which already exhibit many cystic follicles (CF, tertiary follicles) amid least mature follicles. In image **d**, fibrosis is evident on the ovarian surface (arrow).

The fallopian tubes (oviducts) are bilateral, tortuous and tubular structures that extend from the ovary to the uterine horns and are divided into infundibulum, ampulla and isthmus. The fallopian tubes transport the ovum from the ovary and the spermatozoa from the site of deposition to the site of fertilization. The histologic structure is composed of an internal mucosal layer covered by a simple or pseudostratified columnar epithelium with some ciliated cells. The mucosa layer is continuous with the submucosa, consisted of loose connective tissue. The tunica mucosa-submucosa is folded and covered by a thin muscular layer consisting mostly of circular smooth muscle bundles. Externally, the tunica serosa contains many blood vessels and nerves with a superficial mesothelial cell layer and the connective tissue from the mesosalpinx is observed at one pole and is part of the lining of the abdominal cavity representing a fold of the broad ligament that stretches from the ovary to the uterine tube that supports the fallopian tube. Although all oviducts had “normal” histological findings in the control group, nine from the twenty injured oviducts in this series showed mild to moderate fibrosis in the tunica serosa, sometimes associated with a small quantity of mononuclear inflammatory cells (**Figure 5**).

**Figure 5.**
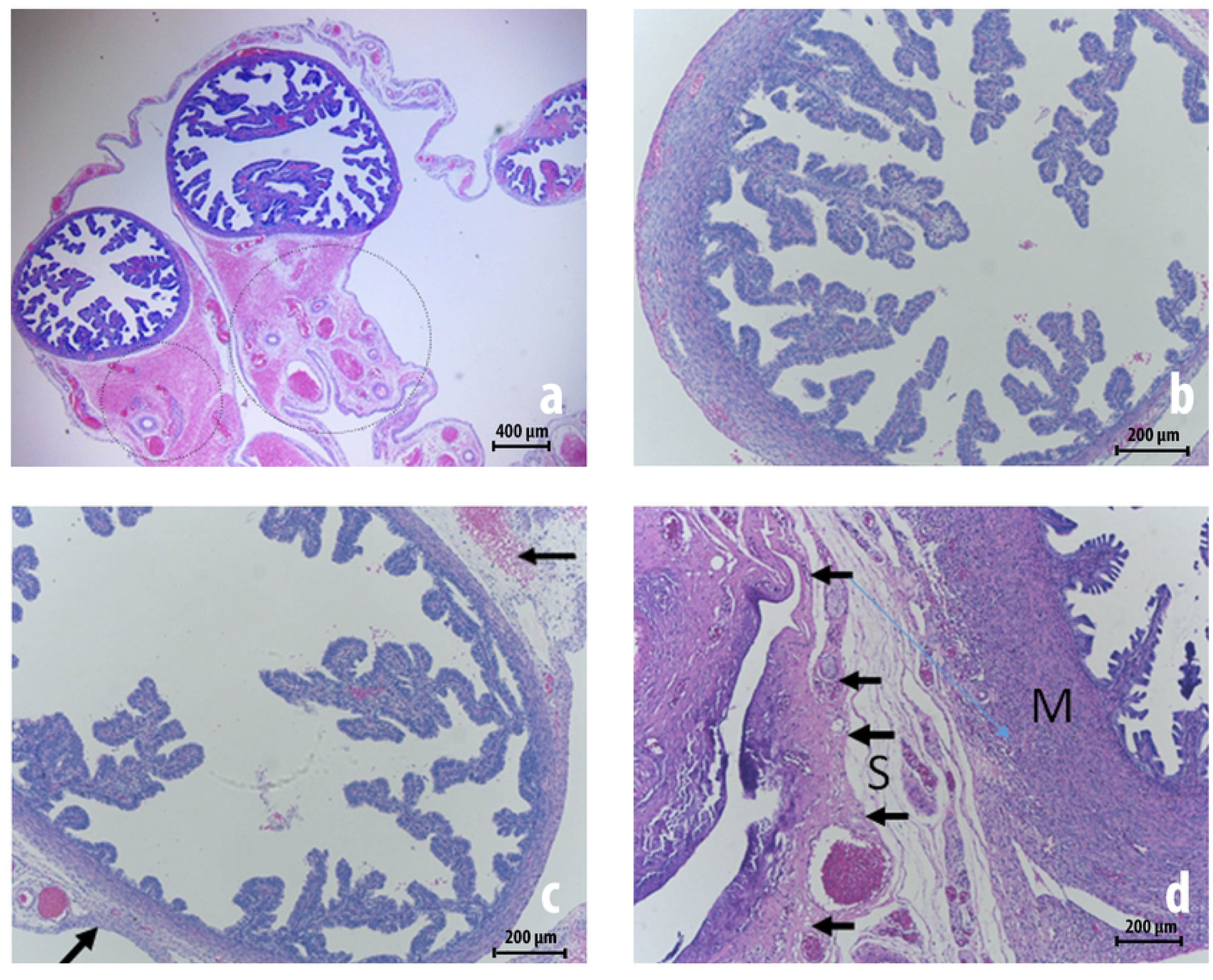
Histology of oviducts prior to and 30 days after peritubal injury (hematoxylin-eosin staining). Images **a** and **b** exhibit micrographs of an oviduct from a 5 ½ months old pig with no previous surgery (control group) showing an internal mucosal layer covered by a simple or pseudostratified columnar epithelium. The tunica mucosa-submucosa is folded and exhibit a papillary architecture – a thin muscular layer (**M**) and, externally, the serosa (**S**) and mesosalpinx with many blood vessels (dotted circles in the image **a**). Images **c** and **d** depict an oviduct from a 6½ month-old pig assessed 30 days after a standardized peritubal laparoscopic injury; there is discrete serositis with edema and inflammatory cells. The arrows point to fibrosis in the serosa.

The uterus in swine has bilateral horns (cornua) connected to the uterine tubes and an unpaired body (corpus). The uterine wall consists of three layers: endometrium with uterine glands covered by pseudostratified columnar epithelium surrounded by a connective tissue (stroma); the myometrium with an thick inner circular smooth muscle and outer longitudinal smooth muscle bundles); and an external layer called serosa (or perimetrium) that is continuous with the corresponding structures in the broad ligament of the uterus. Although all uteri had “normal” histological findings in the control group, some important histopathological findings were observed in the pigs that underwent laparoscopic injury, mainly in the uterine wall (miometrium) and the serosa/perimetrium (**Figure 6**). The main findings consisted of serosa-isolated fibrosis (6 uteri) and fibrosis with inflammatory response associated with giant cell reaction involving a foreign body (attributed to the suture thread) in the myometrium, serosa and the connective tissue from the broad ligament of the uterus (12 uteri). Actually, just one uterine horn was considered histologically normal, that is, with no evidence of local inflammatory response secondary to the prior injury.

**Figure 6.**
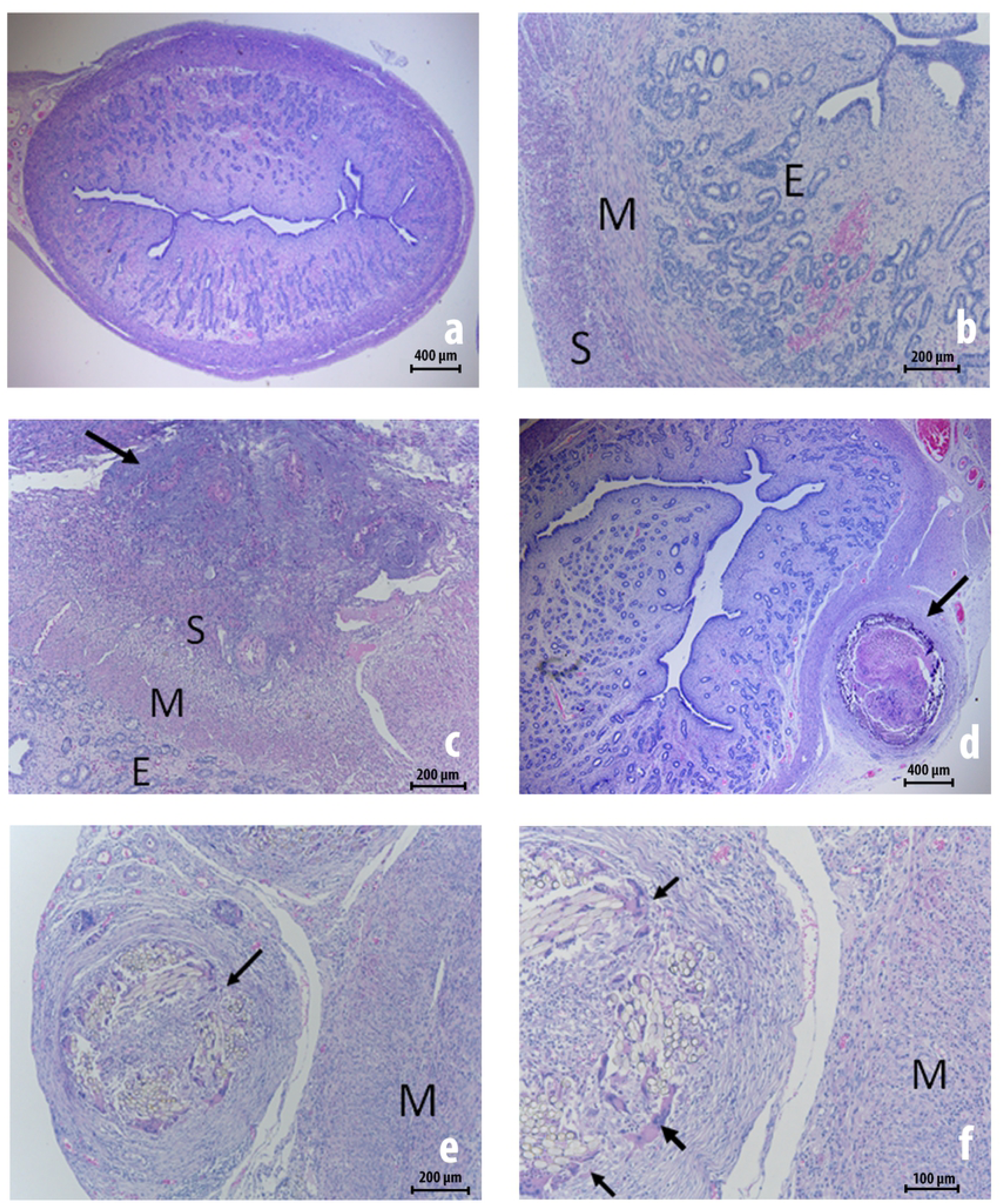
Uterine histology prior to 30 days after peritubal injury (hematoxylin-eosin staining). Images **a** and **b** exhibit micrographs of the uterus of a 5½ month-old surgically naive pig from the control group with normal endometrium (E), myometrium (M) with the inner circular smooth muscle and outer longitudinal smooth muscle bundles, and a thin serosa (S). Images **c, d, e** and **f** exhibit micrographs of the uterus from a 6½ month-old pig assessed 30 days after a standardized peritubal laparoscopic injury; there is fibrosis in the serosa and perimetrium. Giant cell reactions attributed to the suture thread are identified by arrows.

## Biomarkers

The baseline values of blood biomarkers that were assessed immediately before the first surgery (injury) and also immediately before second-look surgery are compared in **Table 2**. Some statistically significant but clinically insignificant changes were noticed for the following assays: serum albumin, globulin, C-reactive protein, fibrinogen, and conjugated and unconjugated bilirubin.

**Table 2.**
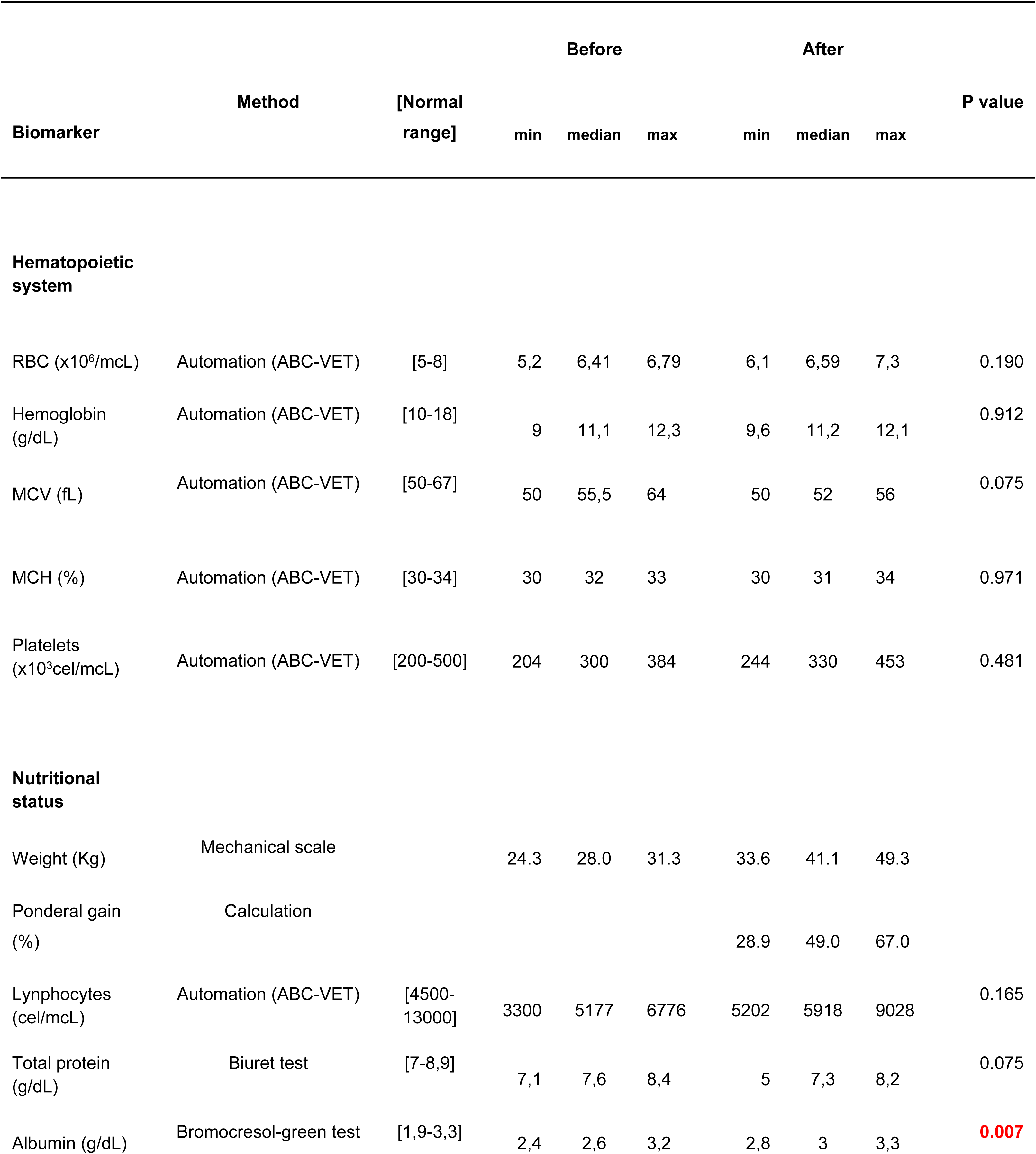
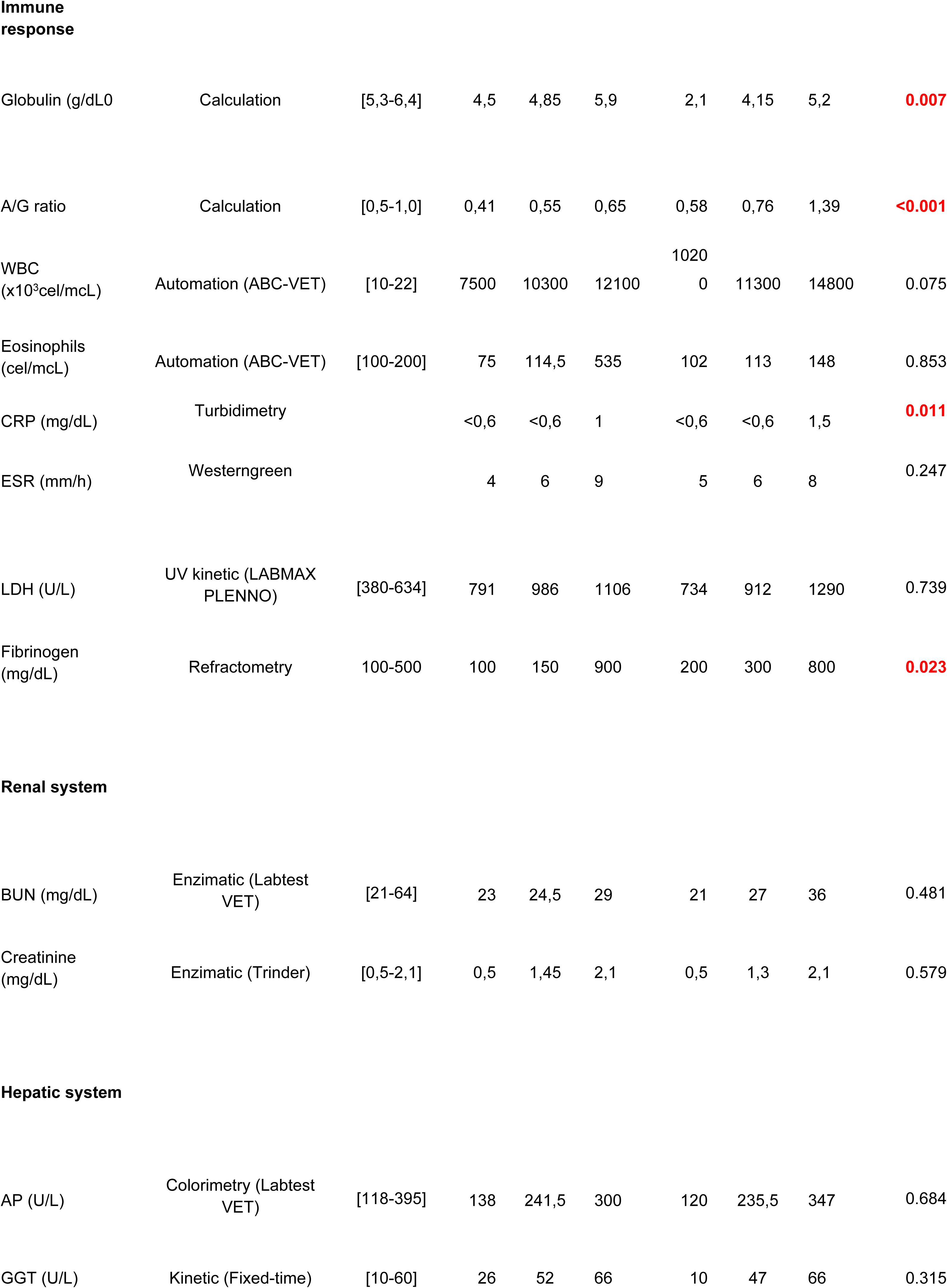
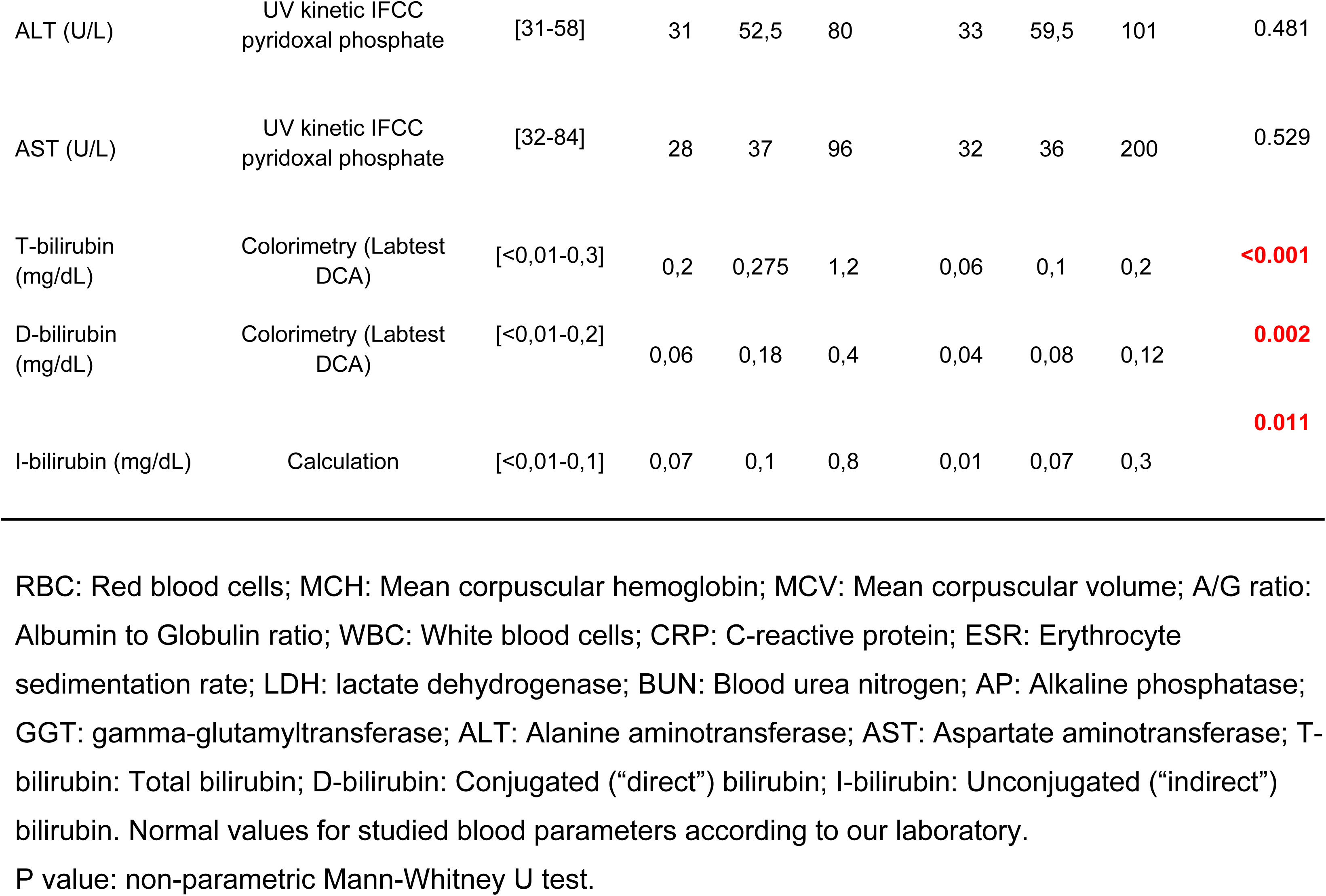
Baseline values of blood biomarkers in 10 young female pigs prior toand 30 days after standardized laparoscopic surgery that was performed to provoke adhesion formation

| Biomarker | Method | [Normal range] | Before |  |  | After |  |  | P value |
| --- | --- | --- | --- | --- | --- | --- | --- | --- | --- |
|  |  |  | min | median | max | min | median | max |  |
| Hematopoietic system |  |  |  |  |  |  |  |  |  |
| RBC (x10<sup>6</sup>/mCL) | Automation (ABC-VET) | [5-8] | 5,2 | 6,41 | 6,79 | 6,1 | 6,59 | 7,3 | 0.190 |
| Hemoglobin (g/dL) | Automation (ABC-VET) | [10-18] | 9 | 11,1 | 12,3 | 9,6 | 11,2 | 12,1 | 0.912 |
| MCV (fL) | Automation (ABC-VET) | [50-67] | 50 | 55,5 | 64 | 50 | 52 | 56 | 0.075 |
| MCH (%) | Automation (ABC-VET) | [30-34] | 30 | 32 | 33 | 30 | 31 | 34 | 0.971 |
| Platelets (x10<sup>3</sup>cel/mCL) | Automation (ABC-VET) | [200-500] | 204 | 300 | 384 | 244 | 330 | 453 | 0.481 |
| Nutritional status |  |  |  |  |  |  |  |  |  |
| Weight (Kg) | Mechanical scale |  | 24.3 | 28.0 | 31.3 | 33.6 | 41.1 | 49.3 |  |
| Ponderal gain (%) | Calculation |  |  |  |  | 28.9 | 49.0 | 67.0 |  |
| Lymphocytes (cel/mCL) | Automation (ABC-VET) | [4500-13000] | 3300 | 5177 | 6776 | 5202 | 5918 | 9028 | 0.165 |
| Total protein (g/dL) | Biuret test | [7-8,9] | 7,1 | 7,6 | 8,4 | 5 | 7,3 | 8,2 | 0.075 |
| Albumin (g/dL) | Bromocresol-green test | [1,9-3,3] | 2,4 | 2,6 | 3,2 | 2,8 | 3 | 3,3 | 0.007 |
| Immune response |  |  |  |  |  |  |  |  |  |
| Globulin (g/dL) | Calculation | [5,3-6,4] | 4,5 | 4,85 | 5,9 | 2,1 | 4,15 | 5,2 | 0.007 |
| A/G ratio | Calculation | [0,5-1,0] | 0,41 | 0,55 | 0,65 | 0,58 | 0,76 | 1,39 | <0.001 |
| WBC (x10<sup>3</sup>cel/mcL) | Automation (ABC-VET) | [10-22] | 7500 | 10300 | 12100 | 1020<br>0 | 11300 | 14800 | 0.075 |
| Eosinophils (cel/mcL) | Automation (ABC-VET) | [100-200] | 75 | 114,5 | 535 | 102 | 113 | 148 | 0.853 |
| CRP (mg/dL) | Turbidimetry |  | <0,6 | <0,6 | 1 | <0,6 | <0,6 | 1,5 | 0.011 |
| ESR (mm/h) | Westerngreen |  | 4 | 6 | 9 | 5 | 6 | 8 | 0.247 |
| LDH (U/L) | UV kinetic (LABMAX PLENNO) | [380-634] | 791 | 986 | 1106 | 734 | 912 | 1290 | 0.739 |
| Fibrinogen (mg/dL) | Refractometry | 100-500 | 100 | 150 | 900 | 200 | 300 | 800 | 0.023 |
| Renal system |  |  |  |  |  |  |  |  |  |
| BUN (mg/dL) | Enzimatic (Labtest VET) | [21-64] | 23 | 24,5 | 29 | 21 | 27 | 36 | 0.481 |
| Creatinine (mg/dL) | Enzimatic (Trinder) | [0,5-2,1] | 0,5 | 1,45 | 2,1 | 0,5 | 1,3 | 2,1 | 0.579 |
| Hepatic system |  |  |  |  |  |  |  |  |  |
| AP (U/L) | Colorimetry (Labtest VET) | [118-395] | 138 | 241,5 | 300 | 120 | 235,5 | 347 | 0.684 |
| GGT (U/L) | Kinetic (Fixed-time) | [10-60] | 26 | 52 | 66 | 10 | 47 | 66 | 0.315 |
| ALT (U/L) | UV kinetic IFCC<br>pyridoxal phosphate | [31-58] | 31 | 52,5 | 80 | 33 | 59,5 | 101 | 0.481 |
| AST (U/L) | UV kinetic IFCC<br>pyridoxal phosphate | [32-84] | 28 | 37 | 96 | 32 | 36 | 200 | 0.529 |
| T-bilirubin<br>(mg/dL) | Colorimetry (Labtest<br>DCA) | [<0,01-0,3] | 0,2 | 0,275 | 1,2 | 0,06 | 0,1 | 0,2 | **<0.001** |
| D-bilirubin<br>(mg/dL) | Colorimetry (Labtest<br>DCA) | [<0,01-0,2] | 0,06 | 0,18 | 0,4 | 0,04 | 0,08 | 0,12 | **0.002** |
| I-bilirubin (mg/dL) | Calculation | [<0,01-0,1] | 0,07 | 0,1 | 0,8 | 0,01 | 0,07 | 0,3 | **0.011** |

RBC: Red blood cells; MCH: Mean corpuscular hemoglobin; MCV: Mean corpuscular volume; A/G ratio:
Albumin to Globulin ratio; WBC: White blood cells; CRP: C-reactive protein; ESR: Erythrocyte
sedimentation rate; LDH: lactate dehydrogenase; BUN: Blood urea nitrogen; AP: Alkaline phosphatase;
GGT: gamma-glutamyltransferase; ALT: Alanine aminotransferase; AST: Aspartate aminotransferase; T-
bilirubin: Total bilirubin; D-bilirubin: Conjugated ("direct") bilirubin; I-bilirubin: Unconjugated ("indirect")
bilirubin. Normal values for studied blood parameters according to our laboratory.
P value: non-parametric Mann-Whitney U test.

All animals showed trace hemoglobin in the urine (+ to ++ / 4+) and 5 pigs had microscopic hematuria (1 to 5 red blood cells per high-power field of urine sediment). These urinalysis findings may simply be a consequence of transdermal suprapubic aspiration technique used to obtain the urine specimens.

## Discussion

This interdisciplinary experimental study describes a porcine model that can be used to assess postoperative peritoneal adhesions, specifically those involving the female reproductive tract that occur after abdominal and pelvic laparoscopic procedures. Besides detailed specifications of the species, our description includes the preoperative preparation (including premedication and the anesthesia regimen), the laparoscopic surgical technique to provoke adhesions in female reproductive tract, the living conditions before the animals undergo a second-look laparoscopic procedure, as well as the ordinal grade and scale that can be used to categorize and quantify the adhesions that have arisen in the interval since the initial surgery. This study also highlights the potential for using this non-rodent animal model to assess the safety of prophylactic agents with regard to possible toxicological effects on multiple systems.

Here we present several key points that may help researchers plan and conduct experimental trials to test prophylactic agents that prevent or minimize adhesion formation. First, the incidence of postoperative adhesions in this porcine model was comparable to that observed in humans, possibly a consequence of taxonomic proximity [10,18,19,20]. Second, by using a laparoscopic approach to perform the injury and to evaluate adhesion formation in this porcine model, researchers can not only perform the same procedures and administer the same prophylactic agents that are contemplated for use in humans, but also carry out a more thorough toxicological assessment of the hepatic, renal, inflammatory and hematopoietic systems than has been performed in the vast majority of studies to date. Third, in addition to assessing adhesion formation and toxicological biomarkers, this animal model also allows a meticulous microscopic evaluation of the potential consequences of exposing reproductive organs (i.e. uterine horn, ovaries and oviducts) to chemical substances through late histopathological observation.

The histopathological evaluations performed 30 days after the laparoscopic injury – in the absence of any prophylactic agent – did not detect alterations to the reproductive tract other than a natural inflammatory response inherent to the tissue healing process. Indeed, all histopathological observations in this series were considered absolutely compatible with the natural development of the reproductive tract of this breed of pig, which reaches maturity between the fifth and sixth months [26] and normal estrus, when well treated under confinement.

Fourth, there are considerations regarding the animal’s size and weight. A pig whose weight is in the range of 30 to 50 Kg is fairly close to the weight of humans. These means it is possible to assess the same kind of injuries associated with laparoscopic procedures and to test the same strategies to prevent adhesion formation using the same devices, kits, agents and doses. Although there could be several potential advantages to using miniature adult pigs which weigh up to 32 Kg and have a slower rate of growth [26], these advantages should be balanced with the benefits of using young larger pigs [9], which may have a very high growth rate. In this series, for example, the median weight gain in 30 days was 13.4 Kg (min 7.9; max 19.1).

Regarding the discrete (but statistically significant) changes observed in the main serum proteins (albumin and globulin), in two inflammatory biomarkers (C-reactive protein and fibrinogen), and in both conjugated and unconjugated bilirubin (**Table 2**), these findings did not point to any specific clinical interpretation, and there was no case in which the blood tests taken together were suggestive of some subclinical pathological condition. Indeed, all the animals were examined by a senior veterinarian (F.L.F.M.) just prior to the second-look surgery and were deemed healthy. Therefore, the main finding concerning the assessed biomarkers was the perception that some discrete changes in blood tests may occur, probably as a natural physiological response, independent of the use of any specific prophylactic agent.

Another point of discussion may be how long we should wait to reassess the animal. The time for remesotheliazation of the peritoneum (or the bridging adhesion) is thought to be no more than 3 to 5 days [27]. Thus the optimal time to assess postoperative adhesions can consider both the operational challenges and cost of caring for the animals for an extended period versus a preference for a longer waiting period, which may favor the identification of late toxic reactions.

Regarding the macroscopic and microscopic assessment of adhesion formation, researchers can elect to assess it directly by necropsy, rather than by laparoscopy. Indeed, although unconventional as compared with the assessment made in living humans, necropsy may be more practical and less expensive in animal models (not assessed in this study). Still, the advantages of a laparoscopic view should not be underestimated. These include the high resolution and about 20-fold magnification, the option of recording video, and the fact that surgeons are accustomed to surveying adhesions in humans during second-look laparoscopic procedures.

## Limitations and strengths

Our study has limitations, particularly, regarding the number of animals. Only 10 animals were used in the intervention arm of the study (and another six animals as controls) and, consequently, the estimates have wide confidence intervals. Of course, this was a consequence of our efforts to adhere to the guiding principles for more ethical use of animals in testing (including the use of methods that enable researchers to obtain comparable levels of information from fewer animals), in accordance with both Brazilian federal laws and Institutional policies that require us to apply the principles of the 3Rs (replacement, reduction, and refinement) to animal research [17].

The main strengths of this experimental study included: (1) the approval by an institutional committee on the ethical use of animals; (2) it used animals with a body mass of the same order of magnitude as humans; (3) it presented a protocol of anesthesia and postoperative analgesia compatible with that offered to humans; (4) it put the main focus on the reproductive tract (uterine horn), on which adhesions are known to cause infertility problems; (5) it used laparoscopic assessment of the adhesions with about 20 times of magnification, as is usually done in humans; (6) it presented practical and feasible ways to collect samples for a more thorough toxicological evaluation (i.e. including hepatic, renal, inflammatory, and hematopoietic systems) than is performed in the vast majority of studies that have been testing prophylactic agents or products; (7) it used, from the taxonomic perspective, a species more closely related to humans as compared to birds, ruminants, rodents and dogs; (8) it optimized the use of animals by offering the possibility of performing laparoscopic training after the experiment was completed and before euthanasia; and (9) it presented an estimated prevalence of adherence (75%) close to real values verified in several clinical studies in humans [10,18,19,20]. Moreover, unlike a study in which a porcine uterine horn adhesion model just mimics a laparoscopic procedure [14], we present an animal model that uses laparoscopic procedures not only to form adhesions, but also to assess them.

## Recommendations

There is great interest in new adhesion prevention technologies [28]. In order to evaluate the risk of toxic effects of prophylactic agents, we suggest the use of multiple tests using blood and urine specimens, not only to detect changes in the systemic inflammatory response, but also to assess the possibility of clinically important adverse effects on the renal, hepatic and hematopoietic systems. From the perspective of protecting the public’s health by ensuring the safety of pharmaceutical and medical devices, the use of this porcine model provides information that may exceed the minimum requirements of the national and international agencies responsible for the toxicological safety of medical products, including those employed to prevent adhesion formation.

With regard to the anesthesia and analgesia used, we recommend premedication with multiple drugs because the synergistic effect of a combination of drugs optimizes the dissociation of the animal from the environment. Acepromazine and midazolam induce muscle relaxation, ketamine guarantees dissociative analgesia during handling until the moment of inhaled induction of general anesthesia, and atropine promotes blockade of sialorrhea and vagal reflexes, so the animal can be manipulated with both minimal suffering and maximum safety [29].

Although we did not focus on controlling this variable, we recommend that an effort should be made so that study animals receive the same quantity of food rations during the postoperative period. One approach would be to segregate them in individual boxes at least during alimentation. In this way, it may be possible to minimize heterogeneity in the individual weight gain (a potential confounder that ranged from 28.9% to 67.0% in this series) (**Table 2**). If individual feeding is feasible, it might also be possible to administer some drugs orally (i.e. dissolved in small amounts of food) rather than parenterally.

Because urine specimens were collected via transdermal suprapubic aspiration, we considered the presence of scant hemoglobin and the occurrence of microscopic hematuria as normal. Such findings should be expected in future studies if urine specimens are obtained using the same technique.

## Conclusion

The use of a porcine model as shown in this study can be a useful *in vivo* animal platform to test the efficacy and safety of prophylactic agents against postoperative peritubal adhesions. We provide several recommendations in order to both minimize animal suffering and avoid problems during experimental trials.

## Sources of Financial or Material Support

This study was supported by DMC Importação e Exportação de Equipamentos Ltda which provides advanced medical devices and equipments for surgical procedures through its product development centers and manufacturing facilities in São Carlos – SP, Brazil, and by Instituto Crispi de Cirurgias Minimamente Invasivas (Rio de Janeiro – RJ, Brazil).

## Conflicts of Interest of the Investigators

The authors have no conflicts of interest.

## Acknowledgements

The authors thank Research and Education Center for Phototherapy in Health Sciences (NUPEN) for the technical support; Dr. Mauren Lopes and Dr. Marina Filgueiras for the careful anesthesia of the animals; Dr. Leigh J. Passman for reviewing the manuscript; and PJ & Christian (Hightec Corporation) for image consulting.

## Notes

Disclosure statement: The authors declare that they have no conflicts of interest and nothing to disclose.

